# Cryopreservation of *Rosa hybrida* cv. Helmut Schmidt by PVS2 vitrification method using *in vitro* fragmented explants (IFEs)

**DOI:** 10.1101/567255

**Authors:** Safiah Ahmad Mubbarakh, Jasim Udain, Jessica Jayanthi James, Rahmad Zakaria, Sreeramanan Subramaniam

## Abstract

This is the first report on cryopreservation via PVS2 vitrification method on roses using *in vitro* fragmented explants (IFEs) as the starting material. The aim of this study is to optimize the efficient plant recovery and regeneration system for cryopreservation of *Rosa hybrida* cv. Helmut Schmidt using IFEs. Some important parameters have been optimized in this study are the effect of ascorbic acid (0.3 mM) examined separately and in combination at all steps in cryopreservation procedure (preculture, loading, unloading and growth recovery), loading type, loading duration, and PVS2 duration. The highest growth recovery of 43.33% was obtained when 3-4 mm size IFEs precultured on 0.25 M sucrose media supplemented with full-strength MS for one (1) day, followed by loading treatment supplemented with 1.5 M glycerol + 0.4 M sucrose + 5% DMSO + 0.3 mM ascorbic acid for 20 minutes, dehydration with PVS2 solution for 30 minutes and then treated with unloading solution supplemented with 1.2 M sucrose + 0.3 mM ascorbic acid for 20 minutes. This finding implies that long-term storage of *Rosa Hybrida* cv. Helmut Schmidt by PVS2 vitrification method was successful with essential biomolecules.

## INTRODUCTION

Roses are one of the most important ornamental plants in many parts of the world. This is because they bloom beautiful flowers and as such, they are widely used in the horticultural industry. Cultivation of rose plants via tissue culture micropropagation has disadvantages in terms of contamination, somaclonal variations and also due to biotic stresses [1]. Hence, cryogenic storage offers a stable and safe method for long term storage of plant materials while safeguarding genetic stability [2]. Cryopreservation at temperatures of −196°C has been considered as an ideal means for long term storage of plant germplasm. Through cryopreservation, hybrids that are in danger of being extinct can be conserved with low maintenance apart from being easy for international exchange. Cryopreservation through vitrification reduces injury caused by intracellular crystallization and provides high regeneration rates prior to immersion into liquid nitrogen. Basically, the most appropriate plant part utilized is the meristem culture since it generates virus free lines [3]. For PVS2 cryopreservation study in roses, *in vitro* fragmented explants (IFE) represent a new source of explant as starting material. IFEs were first introduced by authors [4] who conducted research on Vanilla planifolia “Andrews”. The study discovered that IFEs allows the production of a larger number of shoots than micro cuttings after 4 months in culture. However, every species of plants has different genetic makeup and respond differently to various treatments. Therefore, improvement of the cryopreservation method is crucial in each step for growth after recovery. Vitrification based methods use cryoprotectants like sucrose, DMSO, PVS2, glycerol, and PVS3 that help removes free water and consequently avoid lethal cryoinjury when immersed in liquid nitrogen. Cryoinjury is the prime reason for viability loss which is due to factors such as excision, dehydration, osmotic stress, and changes in temperature during the cryopreservation method. This method causes the production of reactive oxygen species (ROS) that leads to oxidative damage. ROS such as singlet oxygen, superoxide radical, hydrogen peroxide and hydroxyl radicals are toxic and have the capability of reacting with many biomolecules. They aggravate protein modifications, DNA damage, purine alteration, a β-oxidation lipid which are the by-products of energy-generating processes of photosynthetic and respiratory electron transport chain [5]. In addition, the presence of transition metals such as ferum (present as a component of MS media) creates OH- which is the most reactive chemical species in the world produced as a result of Haber-Weiss mechanism or the Fenton reactions [6]. The antioxidant mechanism that occurs naturally in plants is satisfactory to sustain equilibrium between manufacture and scavenging of ROS, a process generally identified as redox homeostasis. However, due to the plant’s stationary lifestyle, they are endlessly exposed to harsh environmental circumstances that elevate ROS generation [5]. When ROS overpowers the cellular scavenging capacity, it leads to oxidative stress. Ascorbic acid is a water-soluble antioxidant molecule which acts directly to counterbalance superoxide radicals during reductive recycling of the oxidized form of α- tocopherol [7]. It is also becoming evident that ascorbic acid is multipurpose as it plays a vital role in photosynthesis, pathogen defense, redox signaling, growth regulation and metal detoxification [8]. The establishment of a cryopreservation method for a particular species requires step by step optimization starting with a standard protocol [9]. When standard cryopreservation methods are not working for new species, step by step optimization is required because of a lack of knowledge in the outcome of different steps in cryopreservation [3]. Given this scenario, the aim of this study is to establish an appropriate and reproducible PVS2 vitrification method for long term conservation of *Rosa hybrida* cv. Helmut Schmidt and to evaluate the effectiveness of ascorbic acid as an antioxidant.

## MATERIALS AND METHODS

### Plant materials

*In vitro* fragmented explants (IFEs) of *Rosa hybrida* cv. Helmut Schmidt was used as a starting material for the PVS2 vitrification method. For multiplication, *Rosa hybrida* cv. Helmut Schmidt plantlets were subcultured in full-strength MS semi-solid media with 30 g/L sucrose supplemented with 4.0 mg/L BAP + 1.0 mg/L Kinetin + 40 mg/L casein hydrolysate. The cultures were grown in a tissue culture room under 16-hour photoperiod cool fluorescent lamps at 25 ± 2°C (Philips TLD, 36 W) at 150 μmol m^−2^ s^−1^. After 4 weeks, plantlets were separated and dissected aseptically (Plate 1) at the base between 3 to 4 mm in size using an autoclaved graph paper underneath the sterile *Petri dish*.

### Application of ascorbic acid at cryopreservation steps

The effect of ascorbic acid was examined separately and in combination in every critical step of cryopreservation (preculture, loading, unloading, growth recovery). Ascorbic acid adjusted to pH 5.7 was added to all solutions and medium that was autoclaved. Ascorbic acid was added to the medium through filter sterilization method using membrane filters (0.45 μm, Minisart, Sartorius stedim). In this experiment, ascorbic acid was added to all four steps with the removal of ferum from all media and solutions. For the media and solutions without ferum, IFEs were transferred to media containing ferum with the absence of ascorbic acid after 14 days. For the improvement on the effect of ascorbic acid at each step of cryopreservation, 3-4 mm IFEs were chosen from four-week-old cultures which were then precultured in full-strength semi-solid MS media [10] supplemented with 0.25 M sucrose for 1 day at 25°C under 16-hour photoperiod. The precultured IFEs were then transferred to 2 mL sterile cryovials (Nalgene Nunc, United States of America [USA]) and treated with 1.5 mL loading solution containing 2 M glycerol and 0.4 M sucrose [11] at room temperature (28⁰C) for 20 minutes. After loading, IFEs were immersed in 1.5 mL PVS2 solution [12] comprising 30% (w/v) glycerol, 15% (w/v) ethylene glycol, 15% (w/v) DMSO and 0.4 M sucrose in full-strength semi-solid media at 0°C for 20 minutes. Prior to treatment with liquid nitrogen (MVE lab 20, MVE Bio-Medical Division, Chart Industries, Inc., USA), IFEs were replaced with fresh PVS2 solution. Then, the cryovials were immersed into liquid nitrogen (MVE lab 20, MVE Bio-Medical Division, Chart Industries, Inc., USA) at −196°C for 24 hours. Negative control treatment (non-cryopreserved IFEs) refers to IFEs that were not plunged into liquid nitrogen. Within a span of 24 hours, the cryovials were removed from liquid nitrogen and thawed in a water bath for 90 seconds at 40°C. PVS2 solution was withdrawn from the cryovials and substituted with unloading solution containing 1.5 mL of full-strength liquid MS media supplemented with 1.2 M sucrose [13] and incubated at 25°C for 20 minutes. IFEs were then washed with MS basal media to remove excess unloading solution. Subsequently, the cryopreserved and non-cryopreserved IFEs were transferred to sterilised filter paper (Whatman plc, United Kingdom [UK]) above full-strength semi-solid MS media (pH 5.7) with 30 g/L sucrose and 2.75 g/L Gelrite in the absence of growth regulator (regrowth media) for 24 hours at 25°C. Following this, the IFEs were shifted to regrowth media with the absence of filter paper. IFEs were placed in dark condition for the first week followed by the second week in dim light condition and the third-week IFEs were exposed to cool white fluorescent lamps at 25±2°C under 16-hour photoperiod.

### Effects of loading type on post-cryopreservation viability

For the optimization of loading type, IFEs with the size range of 3-4 mm were dissected from four weeks old cultures which were then precultured in full-strength semi-solid MS media [10] supplemented with 0.25 M sucrose (condition 3) for 24 hours at 25°C under 16- hour photoperiod (Philips TLD, 36 W, 150 μmol.m^−2^.s^−1^). The IFEs were then placed in 2 mL sterile cryovials (Nalgene Nunc, United States of America [USA]) and supplemented with 1.5 mL loading solution of different compositions [L1, L2, L3, L4, L5, L6, and L7] (Table 1) for 20 minutes at room temperature. The subsequent steps are as follows: dehydration with PVS2, storage in LN, thawing, unloading, growth recovery, number of replicates and statistical analysis were performed as described in the previous section.

**Table 1:** Different types of loading solution

| Treatment | Loading type | References |
| --- | --- | --- |
| L1 | 2M glycerol + 0.4M sucrose | Nishizawa et al. 1992 |
| L2 | 2M glycerol + 1.4M sucrose | Nishizawa et al. 1992 |
| L3 | 1.5M glycerol + 0.4M sucrose + 5% DMSO | Matsumoto et al. 1995 |
| L4 | 10% DMSO + 0.7M sucrose | Matsumoto et al. 1995 |
| L5 | 0.5M glycerol + 0.3M sucrose + 10% DMSO | Matsumoto et al. 1995 |
| L6 | 25% PVS2 | Sakai et al. 1990 |
| L7 | 60% PVS2 | Sakai et al. 1990 |

### Effects of loading duration on post-cryopreservation viability

For the optimization of loading type, IFEs with the size range of 3-4 mm were dissected from four-week-old cultures which were precultured in full-strength semi-solid MS media [10] supplemented with 0.25 M sucrose for 24 hours at 25°C under 16-hour photoperiod (Philips TLD, 36 W, 150 μmol.m^−2^.s-^1^). The precultured IFEs were then transferred into 2 mL sterile cryovials (Nalgene Nunc, United States of America [USA]) and supplemented with 1.5 mL loading solution supplemented with 1.5M glycerol + 0.4M sucrose + 5% DMSO (L3) with the presence of 0.3 mM ascorbic acid for 0, 5,10, 20 and 30 minutes at room temperature (30°C ± 2°C).

The subsequent steps are as follows: dehydration with PVS2, storage in LN, thawing, unloading, growth recovery, the number of replicates and statistical analysis were determined as mentioned in the previous section.

### The effects of PVS2 duration on post-cryopreservation viability

For the optimisation of loading type, IFEs with the range of 3-4 mm were dissected from four week old cultures which were then precultured in full-strength semi-solid MS media [10] supplemented with 0.25 M sucrose (condition 3) for 24 hours at 25°C under 16-hour photoperiod (Philips TLD, 36 W, 150 μmol.m^−2^.s^−1^). The precultured IFEs were then transferred to 2 mL sterile cryovials (Nalgene Nunc, United States of America [USA]) containing 1.5 mL loading solution supplemented with 1.5 M glycerol + 0.4M sucrose + 5% DMSO [L3] with the presence of 0.3 mM ascorbic acid for 20 minutes at room temperature. The loading solution was removed and 1.5 mL PVS2 solution [13] composed of 30% glycerol, 15% ethylene glycol, 15% DMSO and 0.4 M sucrose supplemented with full strength liquid MS media was added to the cryovial at 0°C for 0, 10, 20, 30 and 40 minutes. IFEs that were dehydrated in 1.5 mL ice-cold PVS2 solution was replaced with fresh 1.5 mL ice-cold PVS2 and were immersed in LN for 24 hours. Control treatment is defined as the IFEs that were subjected to all treatment including the PVS2 incubation but excluding the cryostorage and thawing procedures.

The subsequent steps are as follows: storage in LN, thawing, unloading, growth recovery, the number of replicates and statistical analysis were determined as described in the previous section.

#### Proline content analysis

Proline content was determined using the method of [14]. IFEs (0.1 g) were collected at each stage of the PVS2 vitrification method. Absorbance was taken at A_520nm_ using a (U- 1900 UV-VIS Spectrophotometer 200Vv, 3J0-0003, Hitachi, Japan) and data were compared with proline standard curve.

#### Carbohydrate content analysis

Carbohydrate content was determined according to the Anthrone method by the author [15]. Fresh samples (0.1 g) were collected in every step of the PVS2 vitrification method. Absorbance was recorded at 625 nm using (U-1900 UV-VIS Spectrophotometer 200Vv, 3J0-0003, Hitachi, Japan).

#### Superoxide dismutase (SOD) assay

The SOD assay was determined according to the method of the authors [16, 17] with some modifications. The photo inhibition percentages of the enzymes in this study were analyzed followed by standard methods (Wilde and Yu, 1998; Dojindo-Molecular-Technologies, 2011).

#### Catalase assay

Catalase assay was determined according to the protocol of the authors [18, 17] with some modifications. Enzyme activity was calculated by using the reported authors [19, 20] method.

#### Ascorbate peroxidase assay

The ascorbate peroxidase activity was determined according to the authors [21] method. The reduction of the ascorbate was calculated using an extinction coefficient of 2.8 mM^−1^.cm^−1^ Enzyme activity was calculated using the method of the authors [19, 20] formula.

#### Water content determination

The water content of IFEs of *Rosa hybrida* cv. Helmut Schmidt at each step of cryopreservation was determined by using the high constant temperature oven method [22].

The IFEs water content percentage was calculated using the formula below [22]:

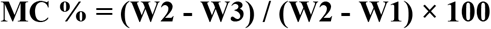

Where,

**W1**= weight of the aluminum boat

**W2**= weight of aluminum boat + IFEs before drying

**W3**= weight of aluminum boat + IFEs after drying

#### Statistical analysis

Regeneration of cryopreserved and non-cryopreserved IFEs was examined 3 weeks after treatment through 2 methods: growth recovery by determining the percentage of regenerated IFEs and the 2,3,5-triphenyl tetrazolium chloride (TTC) spectrophotometrical (U-1900 UV-VIS Spectrophotometer 200Vv, 3J0-0003, Hitachi, Japan) analysis at 490 nm (Verleysen et al. 2004). All treatments consist of six (6) replicates while each treatment comprises of five (5) IFEs. Experiments were performed in a completely randomized design. Means were analyzed through one-way ANOVA and differentiated using Tukey test at p ≤ 0.05. Statistics were performed using IBM SPSS Statistics 22 and results were present as a mean value with standard deviation (SD).

## RESULTS

### Effect of ascorbic acid at different steps of vitrification

Exogenous ascorbic acid added at four important steps of PVS2 vitrification method (Table 2) was utilized by IFEs followed by a significant increase in regrowth of IFEs at some steps. This finding suggests that the cause of death of plant tissues following cryopreservation is due to oxidative stress. Treatment T5 (preculture, loading and unloading steps) and T11 (unloading step) produced 3.33% of regrowth when 0.3 mM ascorbic acid was added at treatment T5 (preculture and loading step) and treatment T1 (preculture step). In addition, treatment T11 and T15 (unloading and growth recovery steps) produced 3.33% and 6.66% regrowth rate after ascorbic acid was added at the unloading and growth recovery step. Post-LN regrowth of shoots from cryopreserved IFEs also significantly improved when 0.3 mM ascorbic acid was added at loading and unloading step (T13) producing regrowth of 20% in terms of growth recovery and also highest TTC value (A490nm: 1.08) (Table 2).

**Table 2:**
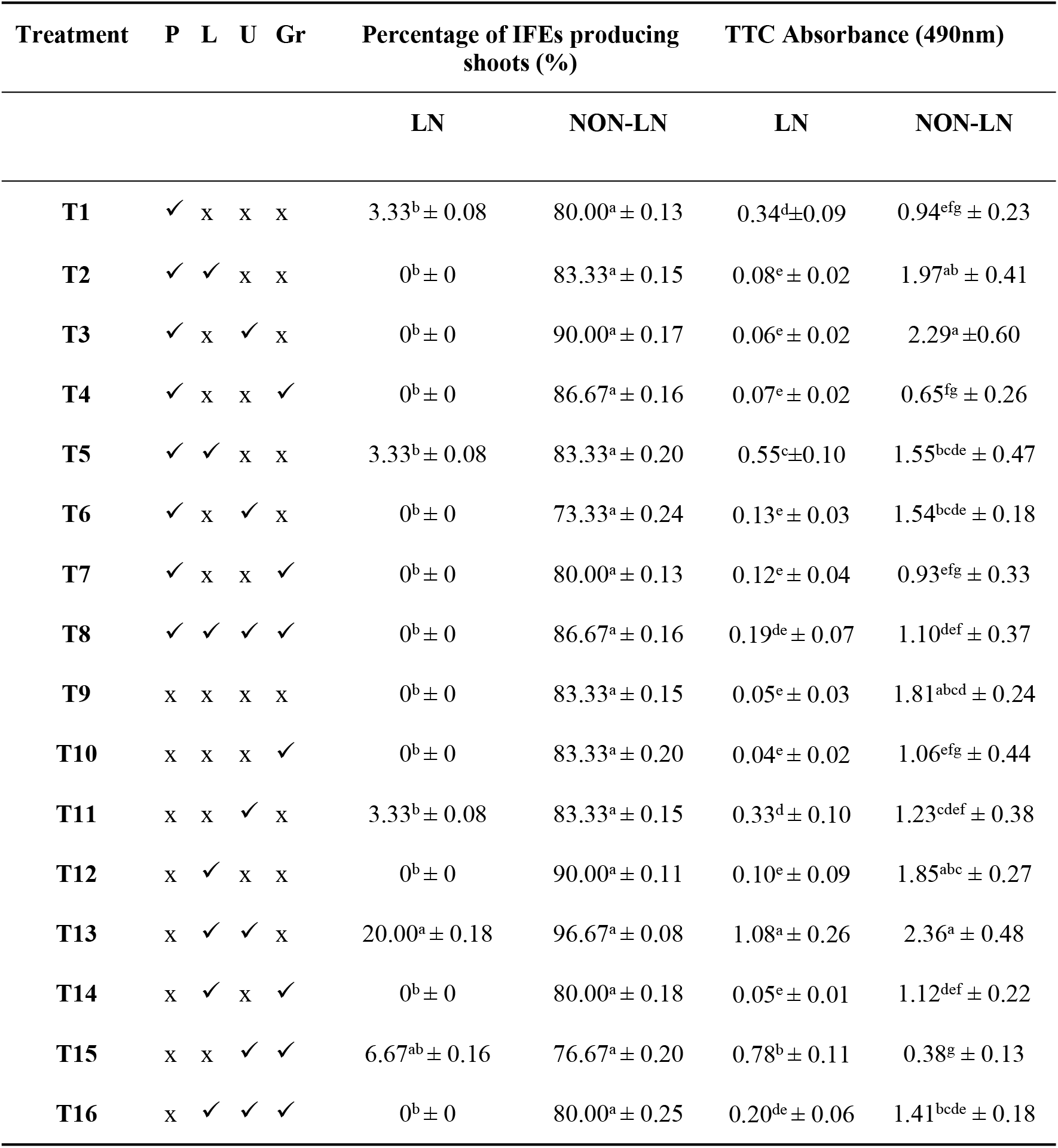
Effect of different combinations of 0.3 mM ascorbic acid on the viability of cryopreserved (LN) and non-cryopreserved (NON-LN) IFEs.

However, supplementation of ascorbic acid in all steps (T8) declined in TTC value and had no sign of regrowth of cryopreserved IFEs. This might be due to the higher concentration of ascorbic acid which could be toxic and damaging to tissues. Similar results were obtained with the absence of ascorbic acid at all steps (T9) but with lower TTC value compared to the addition of ascorbic acid in all steps. Thus, ascorbic acid proves to be an effective treatment for improving the growth recovery of cryopreserved IFEs. Therefore, 0.3 mM ascorbic acid was supplemented at loading and unloading step for the following optimization (a type of loading solutions, loading duration, and PVS2 duration).

The effect of ascorbic acid at different steps in non-cryopreserved IFEs had no significant difference in terms of regrowth percentage and TTC value (Table 2). Supplementation of 0.3 mM ascorbic acid at all steps (T8) produced shoots from IFEs which were decolorized. This confirmed that higher concentrations of ascorbic acid are toxic to the explants. Since treatment T13 (loading and unloading step) gave the highest TTC and regrowth percentage, 0.3 mM ascorbic acid was supplemented at loading and unloading.

### Effect of type of loading solutions on the viability of IFEs

The effect of different compositions of loading solution was examined to study the toxicity levels of solutions and their dehydration capacity towards IFEs tissues. Ascorbic acid (0.3 mM) was supplemented in each loading solution type since the best results obtained from the previous experiments indicated that an addition of 0.3 mM ascorbic acid at loading and unloading step improved the regrowth of IFEs. When viability was determined for cryopreserved IFEs using TTC spectrophotometric assay, the effect of loading type depended on the type of chemical present in the loading solution. Treatment L2 had the least viability (A490nm:0.177) compared to control (L1: 2.0 M glycerol + 0.4 M sucrose) (A490nm: 2.48) and the highest regrowth percentage in general was obtained with the combination of 0.3 mM ascorbic acid + 1.5 M glycerol + 0.4 M sucrose + 5% DMSO (L3) with the percentage of 36.66% growth of shoots from IFEs (A490nm: 2.92). It can be deduced that the chemical compositions of the L3 were ideal both in terms of reduced toxicity level as well as dehydration ability compared to the others which are higher in toxicity causing cell injury. Treatment L1, L4, L5, and L7 produced 20%, 20%, 16.66% and 6.66% growth respectively (Fig 1A). Loading types L2 and L6 failed to produce shoots possibly due to injury caused by chemical toxicity.

**Fig 1.**
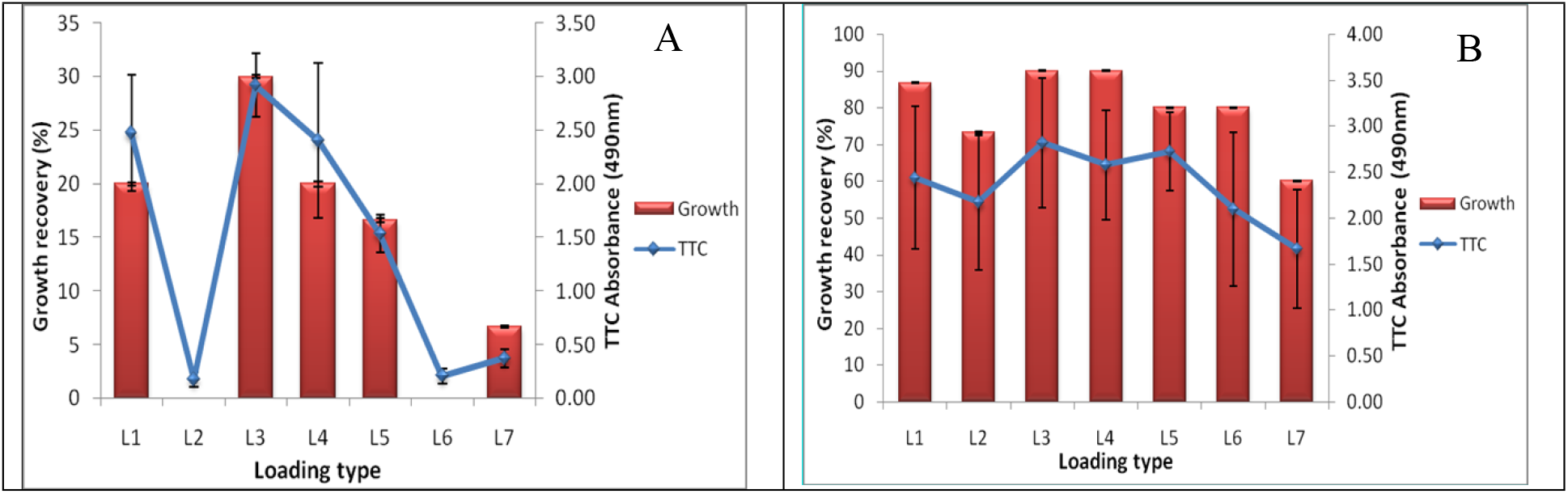
Effect of loading type on the viability of cryopreserved (A) and non-cryopreserved (B) IFEs. Means followed by the same letter were not significantly different using the Tukey test (p≥0.05).

Growth was observed in loading solutions which were supplemented with a higher concentration of DMSO. These results may also be due to the antioxidant impact of DMSO which is a free radical scavenger and may be the reason for the reduced oxidative damage. Therefore in any cryogenic procedure, finding a suitable loading treatment with PVS2 solution involves finding the right balance between toxicity and dehydration. Although there were no significant differences between L1, L3, and L4, L3 which comprise 1.5 M glycerol + 0.4 M sucrose + 5% DMSO was chosen and used for the following experiments. This is because, by utilizing L3 as the loading solution, morphologically healthier shoots were produced from IFEs. Even though DMSO has antioxidant properties it is useful in cryopreservation studies, but higher concentration can be detrimental to plant cells causing genetic changes. Loading solution 3 (L3) [1.5 M glycerol + 0.4 M sucrose + 5% DMSO] was chosen as the loading type for further optimization rather than loading solution 1 (L1) [2 M glycerol + 0.4 M sucrose] which does not contain DMSO and L4 [10% DMSO + 0.7 M sucrose] which contained higher amount of DMSO that may be harmful to plant cells.

Figure 1B shows the effect of the different components of loading solutions on the viability of non-cryopreserved IFEs. The results showed the toxic effect of different types of loading solutions on the viability of non-cryopreserved IFEs without exposure to liquid nitrogen. There were no significant differences observed for types of loading solutions for non-cryopreserved IFEs in terms of TTC value and also regrowth percentage. The highest viability in terms of TTC value and regrowth percentage was the highest with treatment L3 (A490nm: 2.82, 90%) in general. However, the lowest TTC value and regrowth percentage were with L7 (60% of PVS2). This proves that 60 % of PVS2 is very toxic to the tissues of IFEs. Therefore, loading solution L3 which comprise 0.3 mM ascorbic acid + 1.5 M glycerol + 0.4 M sucrose + 5% DMSO was chosen and used for the following experiments.

### The effect of loading durations on the viability of IFEs

Figure 2A shows the effect of loading durations on the viability of cryopreserved IFEs. IFEs were treated with loading solution using 4 different durations (5, 10, 20 and 40 minutes) and with the absence of loading step (0 minutes) which was used as the control in this experiment. Increasing loading duration did not result in increasing IFEs viability. IFEs that were treated with a loading solution for 10 to 20 minutes produced new shoots from IFEs with regrowth of 10.95% and 30% each. This proves that loading treatment is an important step in the cryopreservation process for effective dehydration.

**Fig 2.**
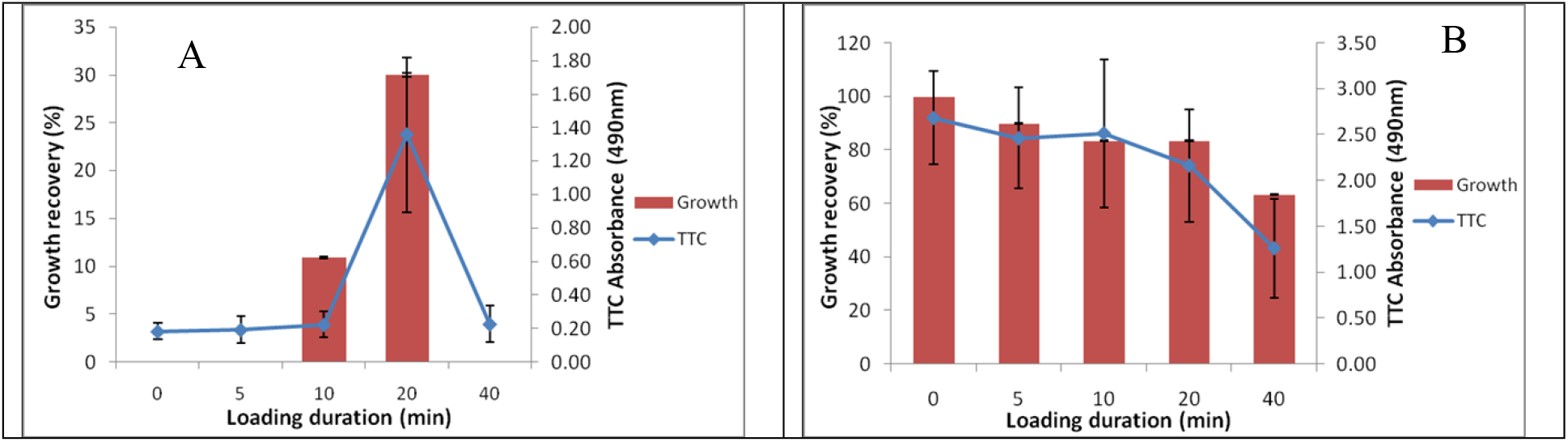
Effect of loading durations on the viability of cryopreserved (A) and non-cryopreserved (B) IFEs. Means followed by the same letter were not significantly different using the Tukey test (p≥0.05).

Figure 2A shows that the greatest survival was achieved with an optimum loading duration at 20 minutes (A490nm:1.36) followed by 10 minutes of treatment (A490nm: 0.224). Therefore, the survival of IFEs was influenced by the loading duration. There was no regrowth on IFEs that were exposed to loading duration for 5 and 40 minutes. Less exposure and extreme exposure to loading solution caused a detrimental effect on IFEs. Thus, the loading duration of 20 minutes was chosen and used for the following experiments.

Figure 2B displayed the effect of loading duration on the survival of non-cryopreserved IFEs. This result displayed the detrimental effect of loading durations which is toxic when IFEs were not subjected to liquid nitrogen. Increasing loading duration causes IFEs viability to decrease gradually. The TTC value and regrowth percentage showed a gradual decrease with the increase in loading duration. The control treatment in this study (0 minutes) which is the absence of a loading step gave the highest TTC value and regrowth percentage (A490nm: 2.69, 100%). There was no significant difference in all the treatments from 5 to 40 minutes. The TTC value decreased gradually (A490nm: 2.46, 2.52, 2.17 and 1.26) after PVS2 treatment at 5, 10, 20 and 40 minutes respectively.

Hence, there was no significant difference for IFEs that were dehydrated for 5, 10 and 20 minutes except for 40 minutes duration when the TTC value was drastically reduced. Regrowth of IFEs was 90% when treated for 5 minutes and significantly decreased to 63.33% after dehydration for 40 minutes.

### The effect of the dehydration period on the viability of IFEs

In the previous experiment, IFEs supplemented with 0.3 mM ascorbic acid + 1.5 M glycerol + 0.4 M sucrose + 5% DMSO (L3) for a duration of 20 minutes produced highest shoots regeneration from IFEs. Therefore, these conditions were further used in PVS2 duration experiment. Figure 3 exhibited the outcome of PVS2 dehydration duration on IFEs survival three weeks post-thaw cryopreservation. IFEs were dehydrated using PVS2 solution using 4 different durations (10, 20, 30 and 40 minutes) and with the absence of dehydration (0 minutes) which was used as the control in this experiment. All cryopreserved IFEs with the absence of dehydration with PVS2 solution (0 minutes) showed very low TTC value (A490nm: 0.17) with no regrowth of shoots. This proves that PVS2 dehydration plays an important role in cryopreservation studies.

**Fig 3.**
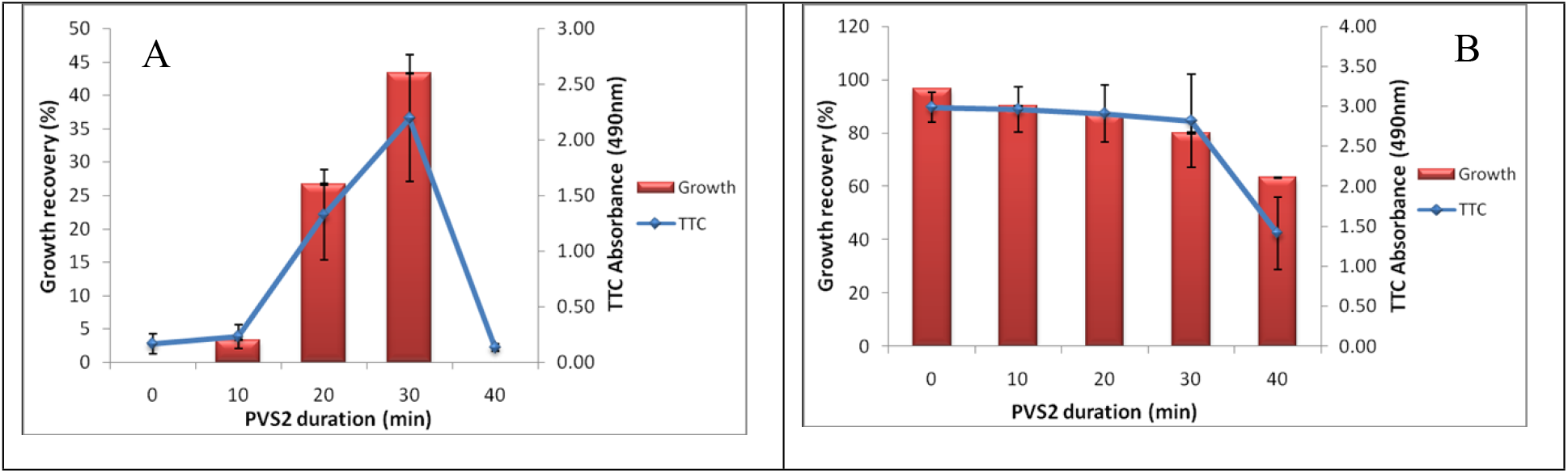
Effect of PVS2 duration on the viability of cryopreserved (A) and non-cryopreserved (B) IFEs. Means followed by the same letter were not significantly different using Tukey test (p≥0.05).

Cryopreserved IFEs (Fig 3A) showed a general increase in TTC value and regrowth percentage after dehydrated in PVS2 solution for 10 minutes and 20 minutes respectively compared to the control IFEs. The regrowth (43.33%) and TTC value (A49nm: 2.20) were significantly highest when PVS2 exposure duration lasted 30 minutes. This shows that the optimum duration to PVS2 solution contributes to successful regeneration. In contrary, TTC value (A490nm: 0.13) reduced drastically with no regrowth of new shoots from IFEs after dehydrated for a duration longer than 30 minutes. In this study, lengthened PVS2 duration displayed deterioration on IFEs survival rate.

Figure 3B demonstrated the effect of PVS2 dehydration duration on the survival of non-cryopreserved IFEs. In the absence of liquid nitrogen, a PVS2 solution showed toxic effects on IFEs. The TTC value and regrowth percentage showed a gradual decrease with the increase in PVS2 duration. The control in this study (0 minutes) refers to IFEs that were not treated with PVS2 solution (absence of dehydration). The control gave produced highest TTC value (A490nm: 2.99) and regrowth percentage (96.67%). The TTC value dropped to A490nm: 2.96, 2.91, 2.82 and 1.42 after PVS2 treatment at 10, 20, 30 and 40 minutes respectively.

There was no significant difference for IFEs that were dehydrated for 10, 20 and 30 minutes except for 40 minutes duration when the TTC value and also regrowth percentage drastically reduced. Regrowth of IFEs was 90% when treated for 10 minutes and significantly decreased to 63.33% after dehydration for 40 minutes.

The optimized method for *Rosa hybrida* cv. Helmut Schmidt was IFEs with size 3-4 mm was precultured in 0.25M sucrose for 24 hours followed by a loading treatment with 1.5 M glycerol + 0.4 M sucrose + 5% DMSO + 0.3 mM ascorbic acid for 20 minutes. After loading, IFEs were treated with PVS2 solution for 30 minutes followed by unloading treatment with supplementation of 0.3 mM ascorbic acid for 20 minutes (Fig 3).

#### Biochemical analysis

#### Proline content

Proline concentration from each cryopreservation stage was estimated based on proline standard curve. The highest amount of proline content for cryopreserved IFEs was observed at thawing (87.05 nmol/mg) and unloading (83.35 nmol/mg) stages when compared to control treatment. However, after 3 weeks of culture in growth recovery media (recovery 2), the proline accumulation reduced to 6.52 nmol/mg with no significant difference to the control IFEs (Fig 4). Loading, PVS2, and growth recovery 1 stage had no significant difference with one another (Fig 4).

**Fig 4.**
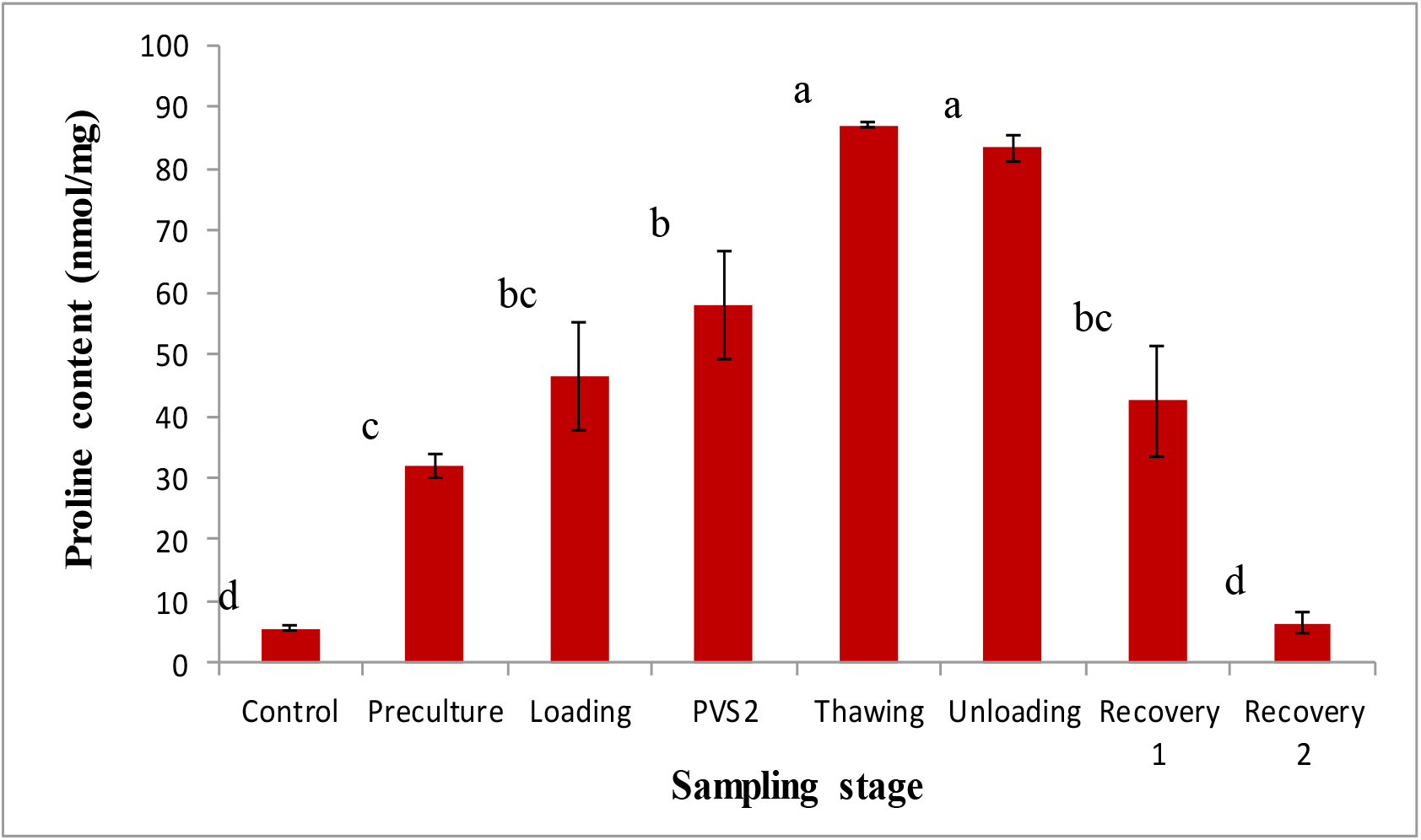
Proline content of cryopreserved IFEs sampled at various stages of the PVS2 cryopreservation method. Results were analyzed using one-way ANOVA. Means followed by the same letter were not significantly different using Tukey test (p≥0.05).

#### Carbohydrate Content

Carbohydrates in the form of soluble sugars, starch and cellulose serve as energy storage and components of cell wall respectively. Carbohydrate content in each cryopreservation stage was determined based on the glucose standard curve.

For cryopreserved IFEs, the highest amount of carbohydrate content (1.757 mg/g) was observed at the unloading stage (Fig 5). However, after 1 day in growth recovery medium (recovery 1) (0.528 mg/g), there was a drastic decrease in carbohydrate content (Figure 5). Recovery 1 stage (0.528 mg/g) had no significant difference with PVS2 (0.632 mg/g) and thawing (0.525 mg/g) stages. Nevertheless, preculture and growth recovery 2 stages showed no significant difference with the control (0.251 mg/g) (Fig 5).

**Fig 5.**
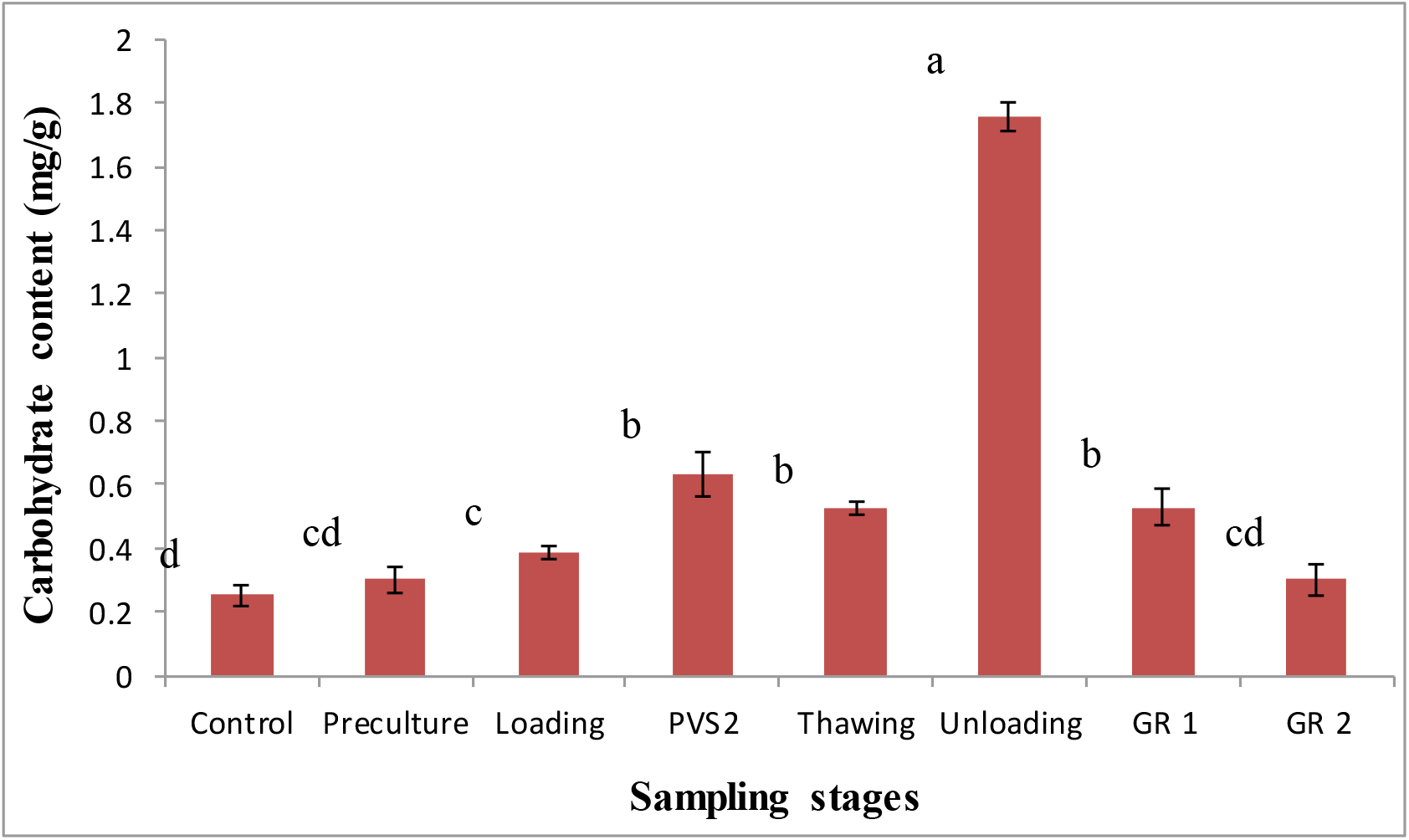
Carbohydrate content of cryopreserved IFEs sampled at various stages of the PVS2 vitrification method. Results were analyzed using one-way ANOVA. Means followed by the same letter were not significantly different using Tukey test (p≥0.05). (GR 1: recovery stage 1, GR 2: recovery stage 2)

#### Superoxide dismutase (SOD)

SOD activity increased during the cryopreserved stages (Fig 6) from the preculture to growth recovery stage. There was an increase in SOD activity at the preculture stage (1516.015 U.g^−1^) with no significant difference to growth recovery stage 1 (1432.950 U.g^−1^). The highest SOD activity was observed at the thawing stage followed by PVS2 stage. The least SOD activity was observed at unloading and growth recovery stage 2 (1168.736 U.g^−1^) (Fig 6).

**Fig 6.**
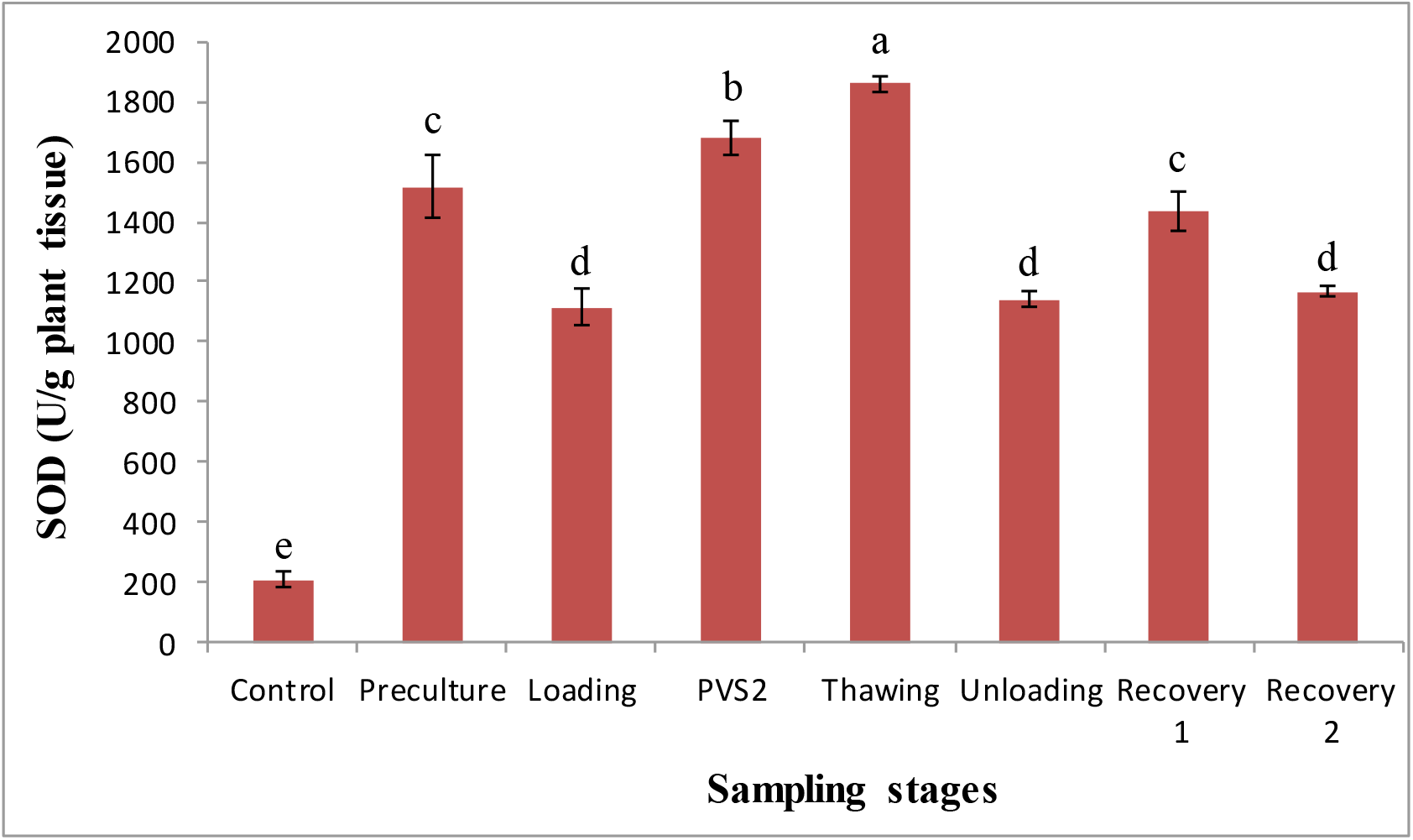
SOD content of cryopreserved IFEs sampled at various stages of the PVS2 vitrification method. Results were analyzed using one-way ANOVA. Means followed by the same letter were not significantly different using the Tukey test (p≥0.05).

#### Catalase (CAT) activity

Cryopreservation procedures significantly affected the catalase activity of cryopreserved IFEs (Fig 7). For cryopreserved IFEs (Fig 7), the highest catalase activity was observed at growth recovery stage 1 (1259.06 U.g^−1^) followed by thawing (1053.454 U.g^−1^) (Fig 7). The catalase activity had no significant difference at preculture (255.475 U.g^−1^), loading (135.3717 U.g^−1^), PVS2 (593.3967 U.g^−1^), unloading (194.4062 U.g^−1^) stages and growth recovery medium (recovery 2) (59.03433 U.g^−1^). All these stages had no significant difference with the control (49.059 U.g^−1^). After a duration of 3 weeks in growth recovery medium (recovery 2), the catalase activity reduced drastically to 59.03433 U.g^−1^.

**Fig 7.**
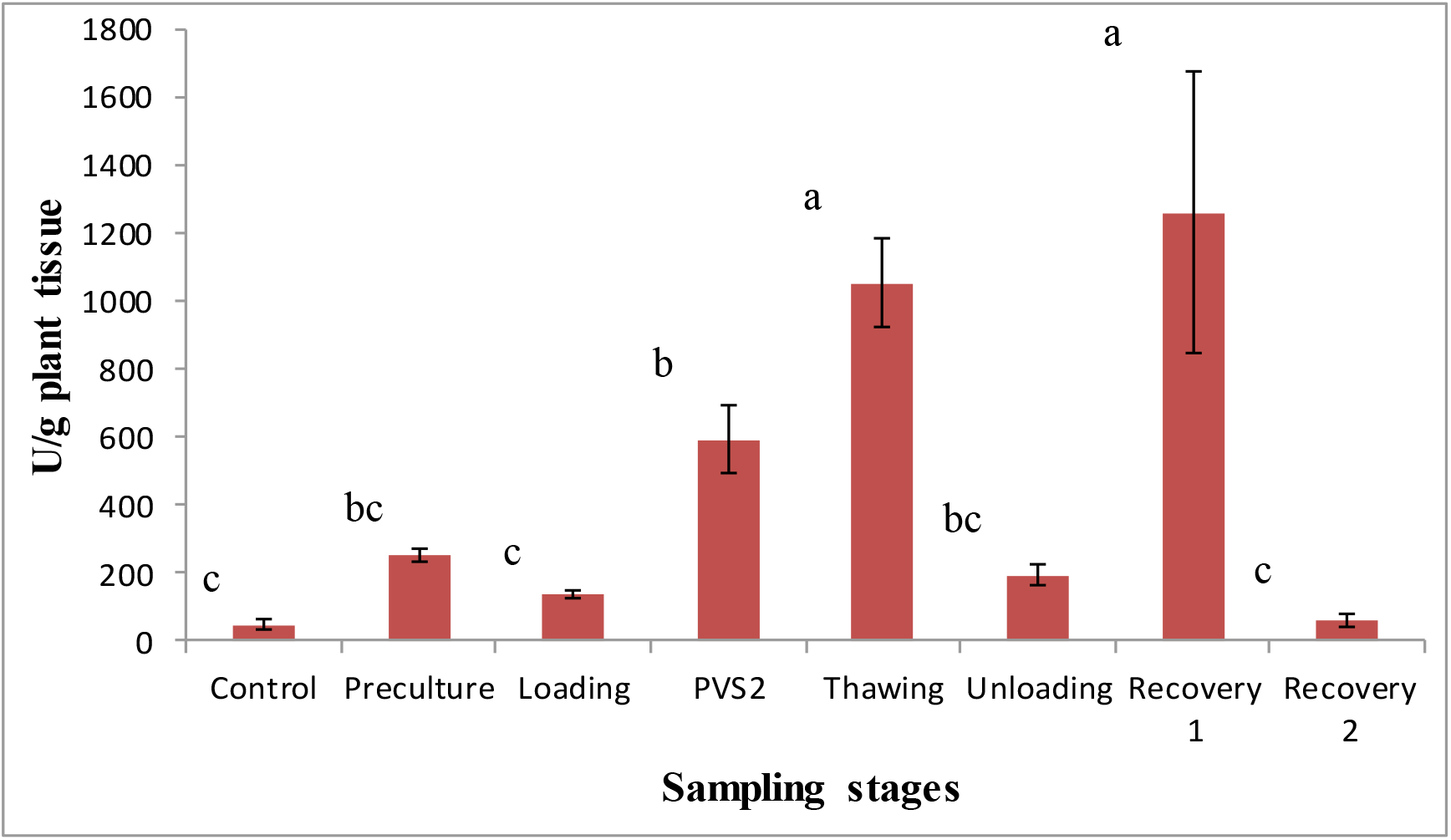
Catalase activity of cryopreserved IFEs sampled at various stages of the PVS2 vitrification method. Results were analyzed using one-way ANOVA. Means followed by the same letter were not significantly different using the Tukey test (p≥0.05).

#### Ascorbate peroxidase (APX) activity

Cryopreservation procedures significantly affected the ascorbate peroxidase activity of treated IFEs (Fig 8). For cryopreserved IFEs (Fig 8), ascorbate peroxidase activity was significantly highest at the thawing stage (491.63 U.g^−1^) followed by PVS2 (259.7 U.g^−1^) compared to low ascorbate peroxidase activity in the control treatment (10.69 U.g^−1^) (Figure 8). The ascorbate peroxidase activity decreased drastically during the unloading (78.07 U.g^− 1^), recovery 1 (34.52 U.g^−1^) and recovery 2 stages (9.57 U.g^−1^). There was a reduction in ascorbate peroxidase activity of cryopreserved IFE post 3-week culture with growth recovery medium (recovery 2) (9.57 U.g^−1^), which was generally lower than the control treatment (10.69 U.mg^−1^) with no significant difference (Fig 8).

**Fig 8.**
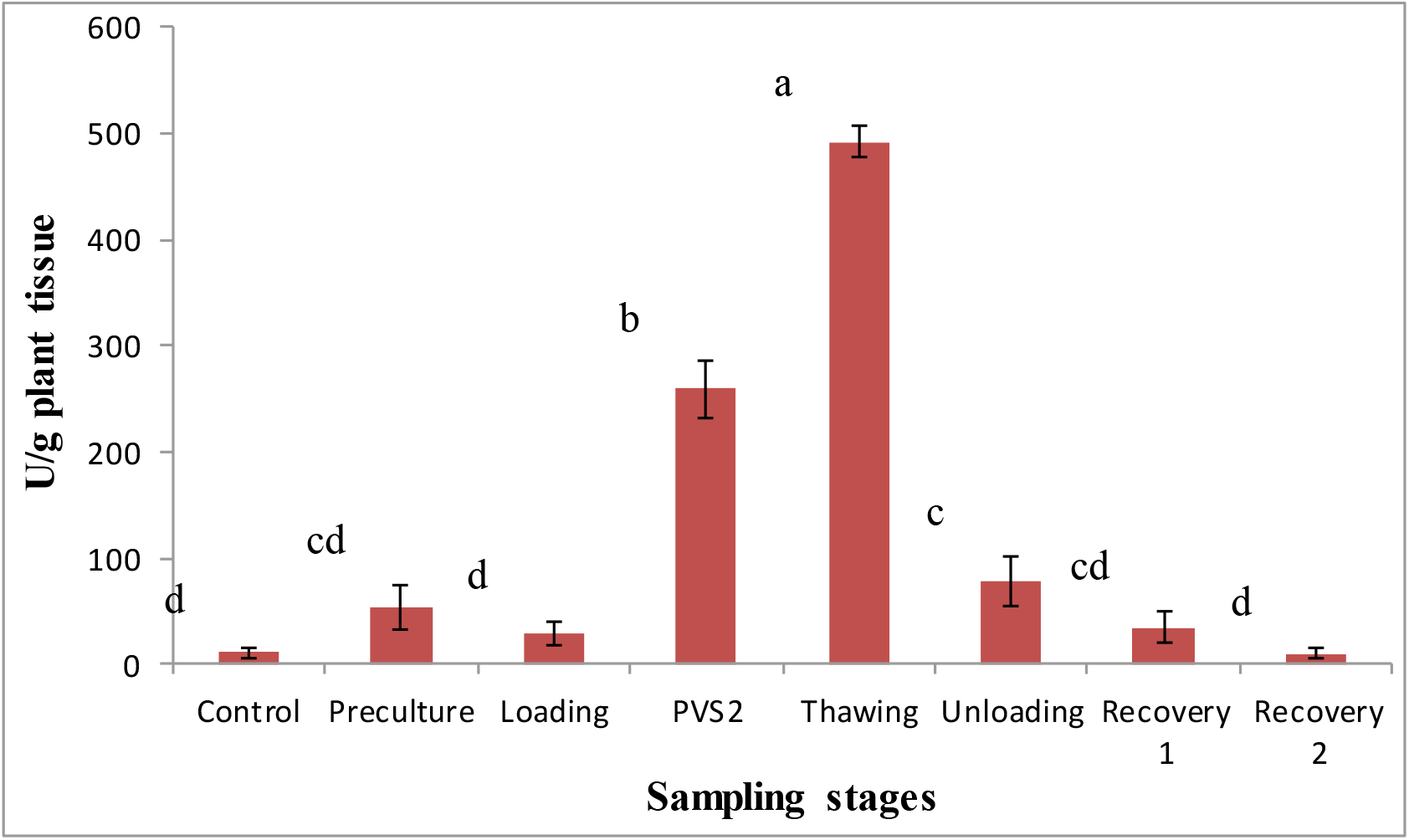
Ascorbate peroxidase activity of cryopreserved IFEs sampled at various stages of the PVS2 vitrification method. Results were analyzed using one-way ANOVA. Means followed by the same letter were not significantly different using the Tukey test (p≥0.05).

#### Water content

The water content of IFEs (Fig 9) was examined to discover the percentage of water lost at each stage of cryopreservation. The water content for cryopreserved IFEs was lowest at the PVS2 dehydration stage (39.21%). Water content which was reduced after control treatment from 93.17% to PVS2 treatment (39.21%) amounted to a total water loss of about 53.96%. After dehydration with PVS2 solution, there was a progressive increase in water content from thawing, unloading, recovery stage 1 and 2 for cryopreserved IFEs (Fig 9).

**Fig 9.**
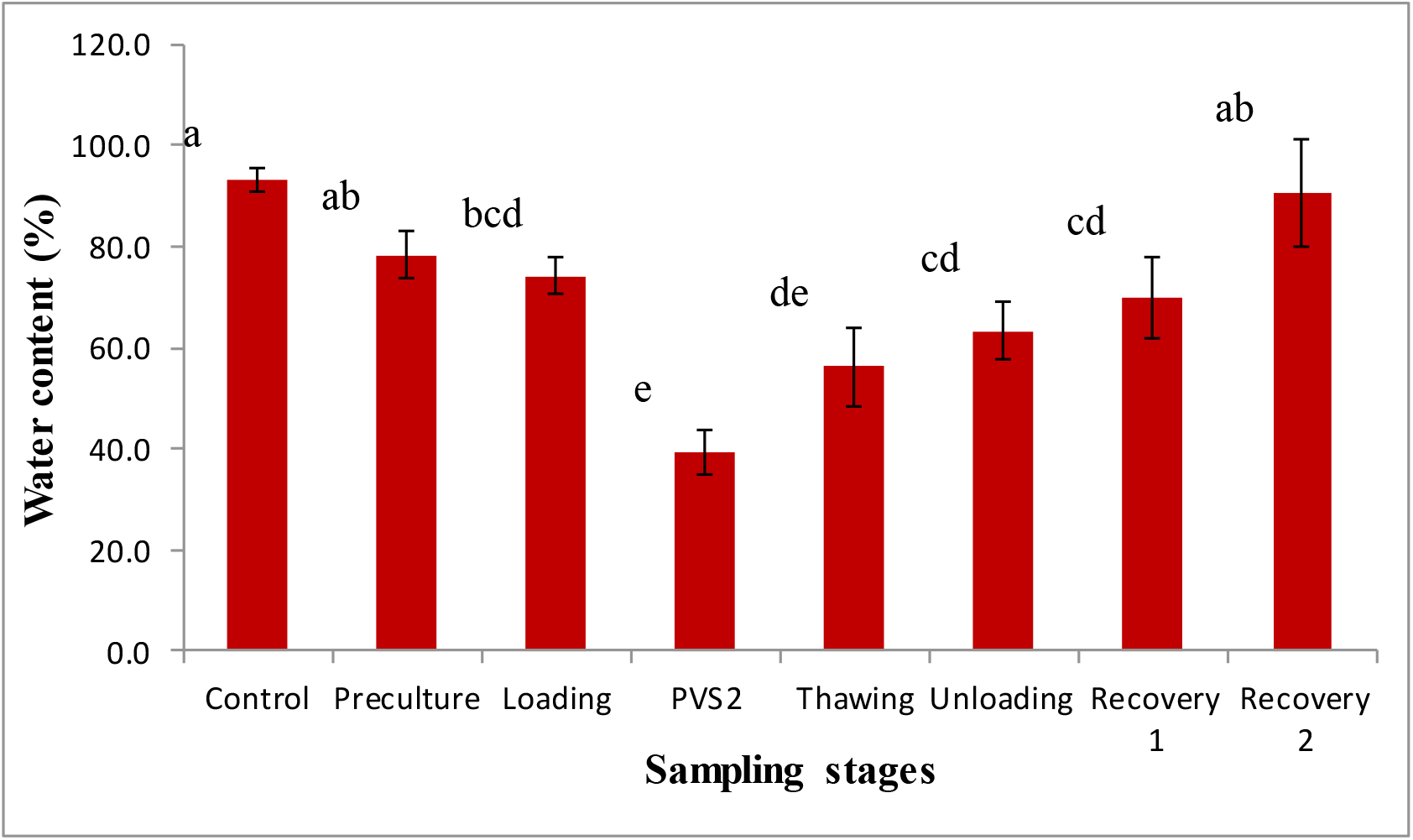
Water content of cryopreserved IFEs sampled at various stages of the PVS2 vitrification method. Results were analyzed using one-way ANOVA. Means followed by the same letter were not significantly different using Tukey test (p≥0.05).

## DISCUSSION

### The effect of ascorbic acid on post-cryopreservation viability

Ascorbic acid (an antioxidant) was included in order to scavenge ROS before oxidative damage occurs. Heavy metals are essential in plant growth and development because they are responsible for influencing cellular processes such as photosynthesis and hemostasis [23]. Most cellular processes require the availability of heavy metals such as Zn, Fe, Cu, and Mn because they are necessary for proper enzyme functioning [24]. Ascorbic acid with the presence of forum participates in Fenton reaction which fabricates the formation of hydroxyl radicals in cells [25]. Ferrum caused growth decline in Arachis hypogaea L. [26] and Blackberry shoot tips [27].

The optimization of the cryopreservation method is important in cryopreservation, but most of the time they are unsuccessful due to oxidative stress caused by ROS as a result of harsh cryopreservation techniques and by cryoprotectants toxicity. According to the authors [28], the outcome of cell death is often caused by the production of ROS. This is injurious to cells due to the chilling and freezing of plants at cold temperatures.

An efficient antioxidant defense system depends on enzymatic (SOD, PRX, CAT) and non-enzymatic antioxidants (salicylic acid, ascorbic acid, glutathione, carotenoids, and tocopherol). Ascorbic acid is a non-enzymatic antioxidant that is excellent in preventing harm caused by ROS in plants. ROS takes place in all tissues of plants but normally higher in photosynthetic cells and meristems. The small antioxidant molecule ascorbic acid helps in the regulation of photosynthesis, senescence and flowering [29]. It also promotes cell proliferation and cell elongation as well as improving plant tolerance to abiotic stress [30]. In this study, ascorbic acid improved the growth of cryopreserved IFEs.

According to the authors [27], antioxidants were able to inhibit the formation of ROS before oxidative damage can arise hence improving the longevity of blackberry shoot tips after cryopreservation. When the ROS production in cells is higher than the natural antioxidants produced in plant cells, then oxidative damage may occur [27]. Therefore, ascorbic acid added at important steps in this study significantly improved regrowth of shoot from IFEs post-cryopreservation. This study proves that oxidative stress eradication is an important factor in plant cryopreservation studies. The highest growth was achieved when ascorbic acid was supplemented at the loading and unloading step. This hypothesis was proven by the authors [31] after they confirmed that the application of ascorbic acid to shoot tips of Nephelium rambutan-ake’s was helpful in the post-cryopreservation.

However, when ascorbic acid was added at all steps (preculture, loading, unloading and growth recovery), TTC viability decreased producing discolored IFEs after 3 weeks of culture. This finding is in line with the report of authors [27] that an excessive amount of ascorbic acid is damaging to plant cells. Therefore, finding the right combination for the supplementation of ascorbic acid is crucial. In contrast to the study on blackberry shoot tips by the authors [27], ascorbic acid was added at all likely combination during preculture, loading, unloading and growth recovery steps of cryopreservation in the study. This study revealed that ROS played a part in the death of plant tissues during cryopreservation and the application of ascorbic acid has boosted IFEs growth.

In this study, ascorbic acid is not only required at loading step but also at the unloading step as it helps to improve survival pre and post cryogenic storage. Adding ascorbic acid in unloading solution is important because thawing at 40°C influences the regrowth of ice crystals which effects regeneration of shoots. This is due to the transformation from a vitreous to a crystalline stage during rewarming of the PVS2 solution which results in a detrimental effect [32]. Besides ascorbic acid, DMSO (A constituent of the PVS2 solution) is a renowned antioxidant and free radical scavenger that helps to control oxidative destruction during some cryopreservation steps [33]. Therefore, in this study, ascorbic acid was not included in PVS2 solution because it already contains DMSO as one of its constituents.

### The effect of loading treatment on post-cryopreservation viability

The advantages of loading solution treatment in cryopreservation methods were proclaimed for countless plant species [34]. According to the authors [35], loading treatment helped to decrease harmful membrane changes developing from serious dehydration with PVS2. The authors [36] added that the loading solution with an increased amount of sucrose concentrations was reported to increase the survival process of cryopreserved orchid seeds. The author [3] claims that this could be associated with the treating of cells with a loading solution which increases cells osmolarity to overcome osmotic injury caused by vitrification solution.

Tissues were first exposed to loading solution which is formulated by a single or a combination of chemicals. In terms of concentration viscosity, they are much reduced compared to the vitrification solution [4]. Studies have shown that the mixture of different chemicals and concentration of chemicals in vitrification solution yield different outcomes. Thus, the type of chemical used in loading solutions must be able to inhibit the detrimental effects of extreme osmotic stress or chemical toxicity.

This study investigated the different combinations of loading solution on the viability of IFEs after cryopreservation. Loading solution L3 which contained a combination of 0.3 mM ascorbic acid + 1.5 M glycerol + 0.4 M sucrose + 5% DMSO showed the best viability compared to other loading solutions. This shows that the chemical composition of loading solution L3 was ideal both in terms of dehydration capability and toxicity level. Loading solution L2 (2 M glycerol + 1.4 M sucrose) and L6 (25% PVS2) produced the lowest TTC viability. This may be due to loading solution L2 which contains the concentrated composition of osmoticum while L6 contains a lower concentration of osmoticum which did not help in dehydration.

A suitable loading solution is essential for the plant species and explants that are delicate to PVS2. Different combinations of loading solution may give rise to higher survivability of shoots. According to the authors [37], loaded with MS media along with 2.0 M glycerol and 0.4 M sucrose had the capacity to cause dehydration and freezing tolerance in different cells and meristems. However, the authors [38] stated that excellent results were acquired when (10% (w/v) DMSO + 0.7 M sucrose) was used. This was ideal in terms of both toxicity level and dehydration efficiency although 2.0 M glycerol and 0.4 M sucrose is stated as the most frequently used solution.

Besides optimizing loading type, loading duration also plays an important role prior to exposure to concentrated vitrification solution (PVS2). Direct exposure of explants to vitrification solution causes detrimental effects towards plants due to vitrification solution toxicity and osmotic pressure [39]. Therefore, success in cryopreservation is only achieved by loading treatment that induces osmotolerance before exposure to PVS2 solution [40]. In this study, when IFEs were not subjected to loading treatment the survival was the lowest. According to the author [3], the loading step is essential in the cryopreservation method since it reduces shock due to osmotic change of solutes and by direct contact to excessively concentrated vitrification solution.

Direct exposure to concentrated vitrification is damaging owing to the fact that chemical toxicity and osmotic stress are limiting factors in vitrification methods [41]. When non-cryopreserved IFEs were loaded with a loading solution it produced higher survivability compared to cryopreserved IFEs. This may be caused by intracellular ice crystal formation post liquid nitrogen exposure. The highest regrowth was achieved when IFEs were loaded for 20 minutes in this study. Similarly, in the study of *Rosa chinensis* ‘Old Blush’, buds that were loaded with a loading solution up to 20 minutes produced the highest regrowth [1].

In contrast, the authors [42] observed that the regrowth crocus calli improved after loading treatment for 30 minutes which was comparable to this study. Embryogenic calli of Anemarrhena asphodeloides Bunge that was treated with a mixture of 2 M glycerol+ 0.4 M sucrose for 30 minutes displayed high regrowth rate (67.8%) with normal plantlets produced after cryopreservation [43]. Even though the accurate mechanism on how the loading treatment induces osmotolerance to PVS2 is uncertain, loading treatment does reduce the osmotic stress and protects the cell membranes from severe dehydration and freezing [44].

### The effect of PVS2 treatment on post-cryopreservation viability

For a successful vitrification method, it is important to optimize the dehydration process in order to provide a balance between dehydration, chemical toxicity and osmotic pressure [45]. Therefore, in any cryopreservation study, exploring the right balance between chemical toxicity and providing enough dehydration for vitrification to take place is required without intracellular freezing to occur during rapid cooling in liquid nitrogen.

However, direct exposure of explants to cryoprotectant solution is detrimental to plant cells due to its chemical toxicity and osmotic stress [39]. To attain successful cryopreservation by vitrification, loading solution should work in tandem with PVS2 solution to reduce the detrimental damage to the plant cells due to osmotic shock caused by cryoprotectants. It has been revealed that during rapid freezing in liquid nitrogen, dehydration of plant cells with PVS2 solution elevates cells ability to achieve vitrified state and avoids the formation of crystals [46]. The level of tolerance to PVS2 solution differs across species due to genotype and water exchange properties that affect osmotic rate [47]. Hence, successful vitrification experiment requires careful management of vitrification solution.

PVS2 is beneficial in vitrification, but overexposure of PVS2 solution is toxic to plants. Therefore, in order for plant cells to avoid chemical toxicity and osmotic stress caused by PVS2 solution, optimization of the dehydration duration is essential in cryopreservation studies [48].

This study displayed positive effects of lengthening the exposure of PVS2 treatment which displayed a more positive outcome in cryopreserved IFEs compared to non-cryopreserved treatment. Non-cryopreserved IFEs displayed a decrease in survival and regrowth when exposure to PVS2 increased. In the present study, when IFEs were not exposed to PVS2 prior to liquid nitrogen storage, the IFEs were damaged producing low TTC value. Similarly, in the study of cryopreserved Phaius tankervilleae seeds, the authors [49] noted zero survival when seeds were exposed to liquid nitrogen without the treatment with PVS2 solution. The decline in survival is an outcome of an osmotic shock to direct exposure to vitrification solutions [42].

Moisture content in plant cells influences viability post-cryopreservation because cells need to have low moisture content to reduce the formation of crystallization and sufficiently high moisture to circumvent dehydration damage [37]. Shoot regrowth with PVS2 solution for *Rosa hybrida* cv. Helmut Schmidt was achieved when IFEs were exposed to PVS2 solution for a duration of 20 minutes and 30 minutes. By the same token, the survival of cryopreserved Dioscorea bulbifera by vitrification methods increased to about 58.9% after 20 minutes of dehydration with PVS2 [50].

Overexposure to vitrification solution to plant cells is detrimental to the viability of the explants. In this study, exposure to PSV2 solution for more than 30 minutes displayed lower survival and zero regrowth post-cryopreservation. Similar results were obtained in cryopreservation study on PLBs of Brassidium Shooting Star orchid by PVS3 vitrification method that longer dehydration duration was not tolerable [51].

In contrast, some plant species can tolerate longer exposure to PVS2 duration. High tolerance to PVS2 solution was observed in avocado embryogenic cultures even though dehydration duration was as long as 120 minutes. Similarly, incubation in PVS2 for 120 minutes produced a 100% survival rate in Quercus [52] and taro plant [53].

However, there are also some plants that survive better with shorter exposure to PVS2. In a study by the authors [54], 10 minutes of PVS2 duration prior to storage in liquid nitrogen showed a higher germination rate. Shorter PVS2 duration helps in survivability of plant by decreasing possible damage caused by chemical toxicity and osmotic stress. The shortest exposure to a vitrification solution was at 1.5 minutes [55]. It can be deduced that the genotype and physiological status of the plants play a role in the sensitivity towards PVS2 solution [1].

#### Proline content

Stress influences proline accumulation in many organisms which include bacteria, protozoa, invertebrates and plants [56]. The authors [57] also confirmed that proline increases in plants exposed to abiotic and biotic stresses. Proline acts as an organic osmolyte that is widely used as a protection material [58]. Accumulation of free proline in many plants is mainly due to the stress of low temperature [59]. Proline content for cryopreserved IFEs of *Rosa hybrida* cv. Helmut Schmidt was the highest at thawing and unloading treatment (after cryostorage) followed by PVS2, loading and stage recovery 1. Similarly, based on the research conducted on *Avena nuda* L. by the authors [59], proline content only increased after exposure to cold stress at −10°C as compared at 1°C. This study showed that a reduced temperature produced greater proline content. Proline functions as a molecular chaperone that helps in stabilizing the structure of proteins and regulates the antioxidant system [60]. Therefore, an increase in free proline content protects plants against stress and providing them with higher tolerance towards cold stress condition.

#### Carbohydrate content

Photosynthesis produces compounds such as carbohydrates had two important roles in plants. Firstly, carbohydrates provide building blocks for plant structural components such as cellulose which is an important element of plant cell walls. Secondly, carbohydrates provide the plant with energy for plant growth and development. Carbohydrate content of cryopreserved and non-cryopreserved IFEs of *Rosa hybrida* cv. Helmut Schmidt was measured as an indication of chilling and dehydration stress encountered during the PVS2 vitrification method. Carbohydrate content of cryopreserved IFEs increased at the unloading stage followed by PVS2, thawing, and growth recovery 1. Increase in carbohydrate content occurred after cold storage in liquid nitrogen. However, the carbohydrate content of cryopreserved IFEs returned to normal levels after 3 weeks in growth recovery medium. It has been reported that in cold storage there is a net breakdown of starch and the accumulation of sucrose from hydrolysis of starch in lily bulbs [61]. Thus, it can be deduced that soluble sugar accumulation might be increased during cold storage. The authors [62] also justified that there is an increase in sugar content during cold acclimation which plays a role in the protection of cells from freezing injury.

Carbohydrates play a part in supplying proline with NADPH and ATP, as a cryoprotectant, osmoregulator, signaling molecules and reactive oxygen species scavengers that have responsibility against chilling stress [63]. Therefore, the increase of soluble sugars during cold stress increases freezing tolerance in plants [64].

#### Superoxide dismutase, catalase, ascorbate peroxidase, and hydrogen peroxide activity

The enzyme superoxide dismutase (SOD) is efficient in catalyzing superoxide molecules, generating oxygen and hydrogen peroxide. Hydrogen peroxide is a type of ROS that has the capability to manufacture hydroxyl radical (OH) via a process called Fenton reaction involving ferrous iron. Thus, catalase is the main enzyme for the decomposition of hydrogen peroxide into water and oxygen. Catalase (CAT) eliminates hydrogen peroxide once converted to highly reactive hydroxyl radicals via Fenton’s reaction. In the absence of catalase enzyme and increase in SOD levels, the formation of hydroxyl radicals increases from the process of Fenton and Haber and Weiss which causes reduced viability. According to the authors [65], although there are many hydrogen peroxide degrading enzymes in plants, catalase is the most unique as it has a very fast turnover rate. Superoxide dismutase, catalase, ascorbate peroxidase and hydrogen peroxide activity of cryopreserved and non-cryopreserved IFEs of *Rosa hybrida* cv. Helmut Schmidt was measured as an indication of oxidative stress encountered during the PVS2 vitrification method. According to the authors [66], the successive steps of cryopreservation bring about ROS-mediated oxidative stress inducing the increases in SOD activity that can elevate stress tolerance in plant cells. SOD activity of cryopreserved IFEs of *Rosa hybrida* cv. Helmut Schmidt amplified at the thawing indicating an increase in the amount and conversion of the superoxide radical to hydrogen peroxide. There was a reduction in SOD activity at the unloading stage which was due to the supplementation of ascorbic acid that assisted in quenching of ROS during the unloading treatment. Ascorbic acid combats ROS creating a monodehydroascorbate (stable free radical) which is short-lived and is able to convert into dehydroascorbate and ascorbic acid initiating the termination of ROS chain [27].

According to the authors [67], owing to reduced levels in CAT activity at the recovery stage, cryopreserved PLBs of *Dendrobium* Sonia-28 experienced excessive stress which led to poor survival. Similarly, higher tolerance in micro-algae cryopreservation was linked to elevated levels of CAT and low SOD activities [68].

According to the authors [69], production of superoxide may be generated when there is an increase in catalase and ascorbate peroxidase activities due to dehydration effects of PVS2 treatment and cryostorage. The authors [70] reported that catalase and ascorbate peroxidase levels in *Brassidium* shooting star orchid displayed the highest antioxidant activity during dehydration and cryostorage treatments.

The increased expression of antioxidant enzymes such as SOD and catalase allow plants to better tolerate cold-induced oxidative injuries [67]. According to the authors [68], tolerant *Ribes* genotype displayed a higher level of antioxidant throughout recovery while sensitive genotype indicated no change in oxidative stress markers. Therefore, the production and protection of plants enzyme are dependent on plant genotype and species.

#### Water content

Cryopreservation of plant tissues is problematic due to the availability of water in plant cells that can be transformed into ice during cooling and rewarming which is capable of causing permanent damage. Reducing water content in plant cells prior to cryopreservation by treatment with cryoprotectants reduces severe injury due to ice crystallization [71]. Water content is known to significantly affect regrowth and development of plants post-cryopreservation [45].

The water content of both cryopreserved and non-cryopreserved IFEs of *Rosa hybrida* cv. Helmut Schmidt reduced during the PVS2 treatment and increased progressively up to the recovery stage. Water content reduction in plant cells due to dehydration initiated by the exposure to PVS2 solution is a crucial step in cryopreservation studies because water content reduction is required for plant survival and recovery. Dehydration reduces the water content in cells and lowered water content reduces physical damages caused by ice crystals upon freezing [72]. *Rosa hybrida* cv. Helmut Schmidt was dehydrated with PVS2 treatment for 0, 10, 20, 30, and 40 minutes. The highest growth was achieved when IFEs were dehydrated with PVS2 solution for 30 minutes producing a percentage of water content at 39.21% (Fig 9.) According to the authors [45, 73], optimizing the adequate exposure time of plants cells to PVS2 solution is an important step in vitrification. This is because it is the key to finding the balance between suitable water content and minimal chemical toxicity with an end goal to have fruitful survival post-cryopreservation. Responses of various species to liquid nitrogen storage post dehydration vary and are attributed to the genotypic variations between species.

Dehydration helps removal of water that might be converted into ice in cells during cryostorage causing damage and plant death eventually. However, the removal of a sufficient amount of water from plant cells is also very crucial. For example, the optimum water content for *Acer platanoides*, a plant species that produces desiccation-tolerant seeds, was at 10–15 %. On the other hand, recalcitrant seeds of *Acer pseudoplatanus*, achieve the best cryopreservation results with slightly higher water content at 15–20% [74]. Therefore, it can be deduced that different plants withstand the different amount of dehydration.

## Conclusion

Ascorbic acid counteracts ROS by boosting antioxidant capability thus improving growth. The addition of 0.3 mM exogenous ascorbic acid at loading and rinsing step was easy and economical to improve growth of *Rosa hybrida* cv. Helmut Schmidt. Selecting a suitable loading solution (0.3 mM ascorbic acid + 1.5 M glycerol + 0.4 M sucrose + 5% DMSO) and PVS2 treatment for 30 minutes was crucial for the survival of samples post cryostorage. This finding implies that long-term storage of *Rosa hybrida* cv. Helmut Schmidt by PVS2 vitrification method was successful with essential biomolecules.

---

**Plate 1:**
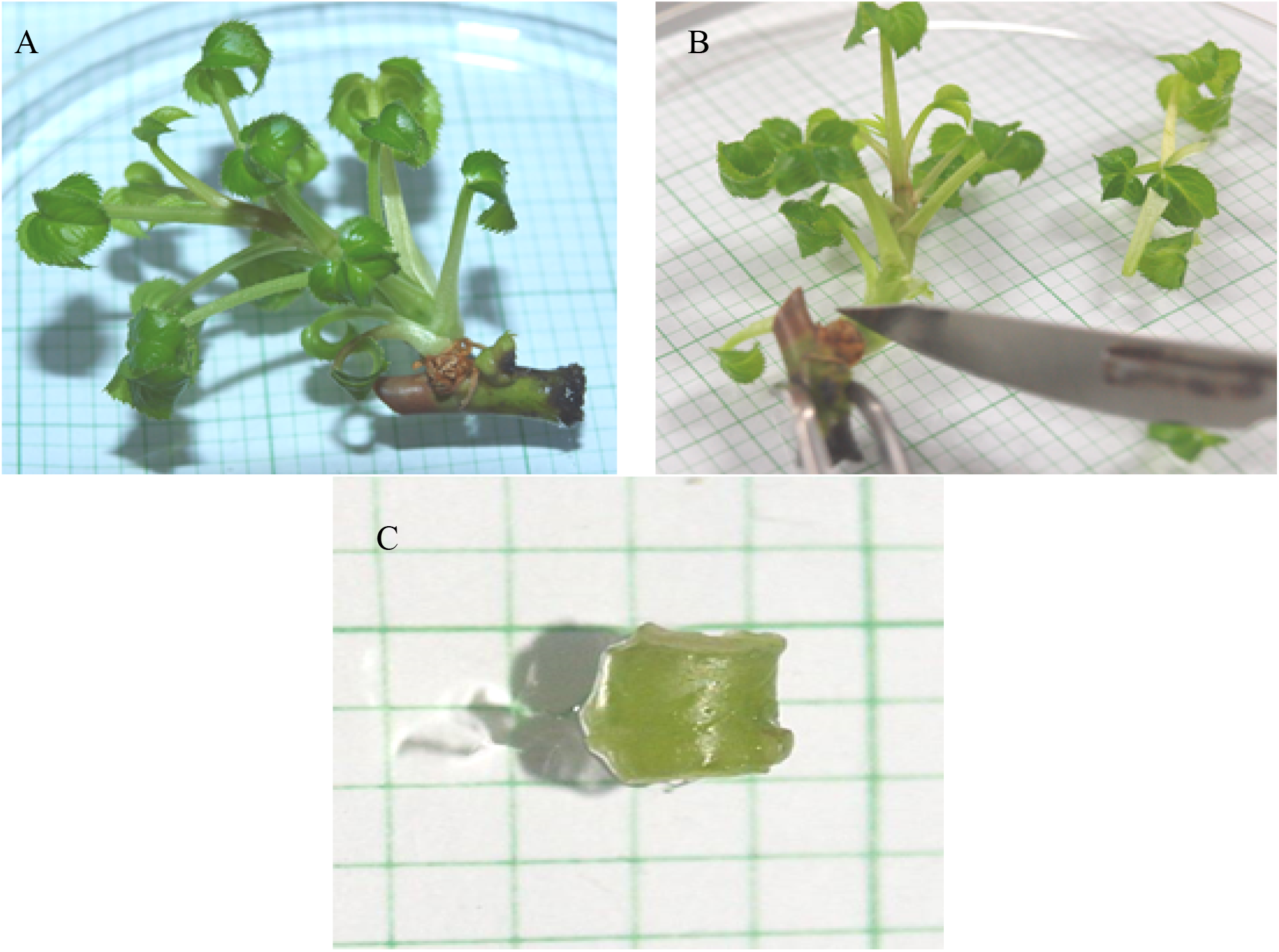
*In vitro* grown *Rosa hybrida* cv. Helmut Schmidt plantlet. (A) Four weeks old *in vitro* plantlets were cleaned to remove oxidized tissues, (B) The base of the *in vitro* plantlets were dissected to about 3 to 4 mm in size, and (C) IFE with size 3 to 4 mm is shown. Scale bar represents 5 mm.

